# Differentiable processing of objects, associations and scenes within the hippocampus

**DOI:** 10.1101/208827

**Authors:** Marshall A. Dalton, Peter Zeidman, Cornelia McCormick, Eleanor A. Maguire

**Affiliations:** Wellcome Centre for Human Neuroimaging, Institute of Neurology, University College London, London WC1N 3AR, UK

## Abstract

The hippocampus is known to be important for a range of cognitive functions including episodic memory, spatial navigation and future-thinking. Wide agreement on the exact nature of its contribution has proved elusive, with some theories emphasising associative processes and another proposing that scene construction is its primary role. To directly compare these accounts of hippocampal function in human males and females, we devised a novel mental imagery paradigm where different tasks were closely matched for associative processing and mental construction, but either did or did not evoke scene representations, and we combined this with high resolution functional MRI. The results were striking in showing that differentiable parts of the hippocampus, along with distinct cortical regions, were recruited for scene construction or non-scene-evoking associative processing. The contrasting patterns of neural engagement could not be accounted for by differences in eye movements, mnemonic processing or the phenomenology of mental imagery. These results inform conceptual debates in the field by showing that the hippocampus does not seem to favour one type of process over another; it is not a story of exclusivity. Rather, there may be different circuits within the hippocampus, each associated with different cortical inputs, which become engaged depending on the nature of the stimuli and the task at hand. Overall, our findings emphasise the importance of considering the hippocampus as a heterogeneous structure, and that a focus on characterising how specific portions of the hippocampus interact with other brain regions may promote a better understanding of its role in cognition.

**Significance statement:** The hippocampus is known to be important for a range of cognitive functions including episodic memory, spatial navigation and future-thinking. Wide agreement on the exact nature of its contribution has proved elusive. Here we used a novel mental imagery paradigm and high resolution fMRI to compare accounts of hippocampal function that emphasise associative processes with a theory that proposes scene construction as a primary role. The results were striking in showing that differentiable parts of the hippocampus, along with distinct cortical regions, were recruited for scene construction or non-scene-evoking associative processing. We conclude that a greater emphasis on characterising how specific portions of the hippocampus interact with other brain regions may promote a better understanding of its role in cognition.

## Introduction

There is long-standing agreement that the hippocampus is essential for supporting memory, especially long-term episodic or autobiographical memory (Scoville and Milner, 1957; Squire, 1992; Clark and Maguire, 2016) and for facilitating spatial navigation (O’Keefe and Nadel, 1978). More recently, the hippocampus has been linked with other roles including scene perception (Graham et al., 2010), short-term memory (Hartley et al., 2007; Hannula and Ranganath, 2008), constructing mental representations of scenes (Maguire and Mullally, 2013; Zeidman and Maguire, 2016), imagining the future (Schacter et al., 2012; Hassabis et al., 2007), decision-making (McCormick et al., 2016; Mullally and Maguire, 2014) and mind-wandering (Karapanagiotidis et al., 2017; McCormick et al., 2018). In addition, accumulating evidence suggests that different hippocampal subfields are preferentially recruited during specific cognitive processes (see examples in Berron et al., 2016; Dimsdale-Zucker et al., 2018; Eldridge et al., 2005; Guzman et al., 2016; Hodgetts et al., 2017; Zeidman et al., 2015a).

Numerous theories attempt to describe how the hippocampus may support such a seemingly diverse range of functions, including the relational theory and scene construction theory. The relational theory proposes that the hippocampus is required for the binding of arbitrary relations among individual elements within an experience or associating items within a context regardless of whether or not these associations are couched within a spatial framework (Konkel and Cohen, 2009;

Cohen and Eichenbaum, 1993). This view has much in common with other theories that place associative processing at the heart of hippocampal function, namely, the binding of item and context model (Ranganath, 2010), the domain dichotomy model (Mayes et al., 2010), the configural theory (Rudy and Sutherland, 1995), the constructive episodic simulation hypothesis (Roberts et al., 2017) and the high resolution binding hypothesis (Yonelinas, 2013).

In contrast, the scene construction theory posits that a prime function of the hippocampus is to construct internal models of the world in the form of spatially coherent scenes. Summerfield et al. (2010) and Mullally and Maguire (2013) found that three objects and a three-dimensional (3D) space are sufficient to form the subjective experience of a scene during mental imagery. This is the operational definition of a scene that we use here. Recently, scene construction has been linked with a specific part of the hippocampus – the anterior medial portion that encompasses the presubiculum and parasubiculum (pre/parasubiculum; Zeidman et al., 2015a; Zeidman et al., 2015b; Zeidman and Maguire, 2016; Hodgetts et al., 2016; Maass et al., 2014; Dalton and Maguire, 2017).

Our goal in the current study was to directly compare the relational/associative and scene construction theories. We devised a novel mental imagery task in which participants engaged in mental construction of objects, non-scene arrays (three objects and a 2D space) and scenes (three objects and a 3D space). These tasks were matched for associative processing but, importantly, only the latter evoked the mental experience of a scene representation. This paradigm, therefore, made it possible to examine whether hippocampal recruitment was modulated by the associative processing that was required for both array and scene construction, or whether the hippocampus was preferentially engaged by scenes. Findings either way would provide novel evidence to inform conceptual debates in the field.

Given previous findings linking the anterior medial hippocampus with scene processing, we predicted that this area would be activated by our scene construction task. We also evaluated the recent relevant prediction, based on anatomical considerations, that the objects task might preferentially activate prosubiculum/CA1 due to direct links with the perirhinal cortex (PRC; Dalton and Maguire, 2017; Insausti and Munoz, 2001). More widely, we predicted that retrosplenial (RSC), posterior cingulate (PCC) and posterior parahippocampal (PHC) cortices would be particularly active during the scene construction task given their known links with scene processing (Epstein, 2008), while the object and array construction tasks would engage PRC, given its acknowledged role in object processing (Nelson et al., 2016; Olarte-Sanchez et al., 2015; Buckley and Gaffan, 1998).

## Materials and Methods

### Participants

Thirty healthy, right handed participants took part in the study (20 females, mean age 24 years, SD 4.12). All had normal or corrected to normal vision and gave written informed consent in accordance with the University College London research ethics committee.

### Tasks and stimuli

The fMRI experiment comprised six separate mental construction tasks: ‘Imagine Fixation’, ‘Imagine Objects’, ‘Imagine 2D Grid’, ‘Imagine 3D Grid’, ‘Construct Array’ and ‘Construct Scene’ (Fig. 1A-F). For each task, participants engaged in mental construction with their eyes open while looking at a blank white screen.

**Figure 1.**
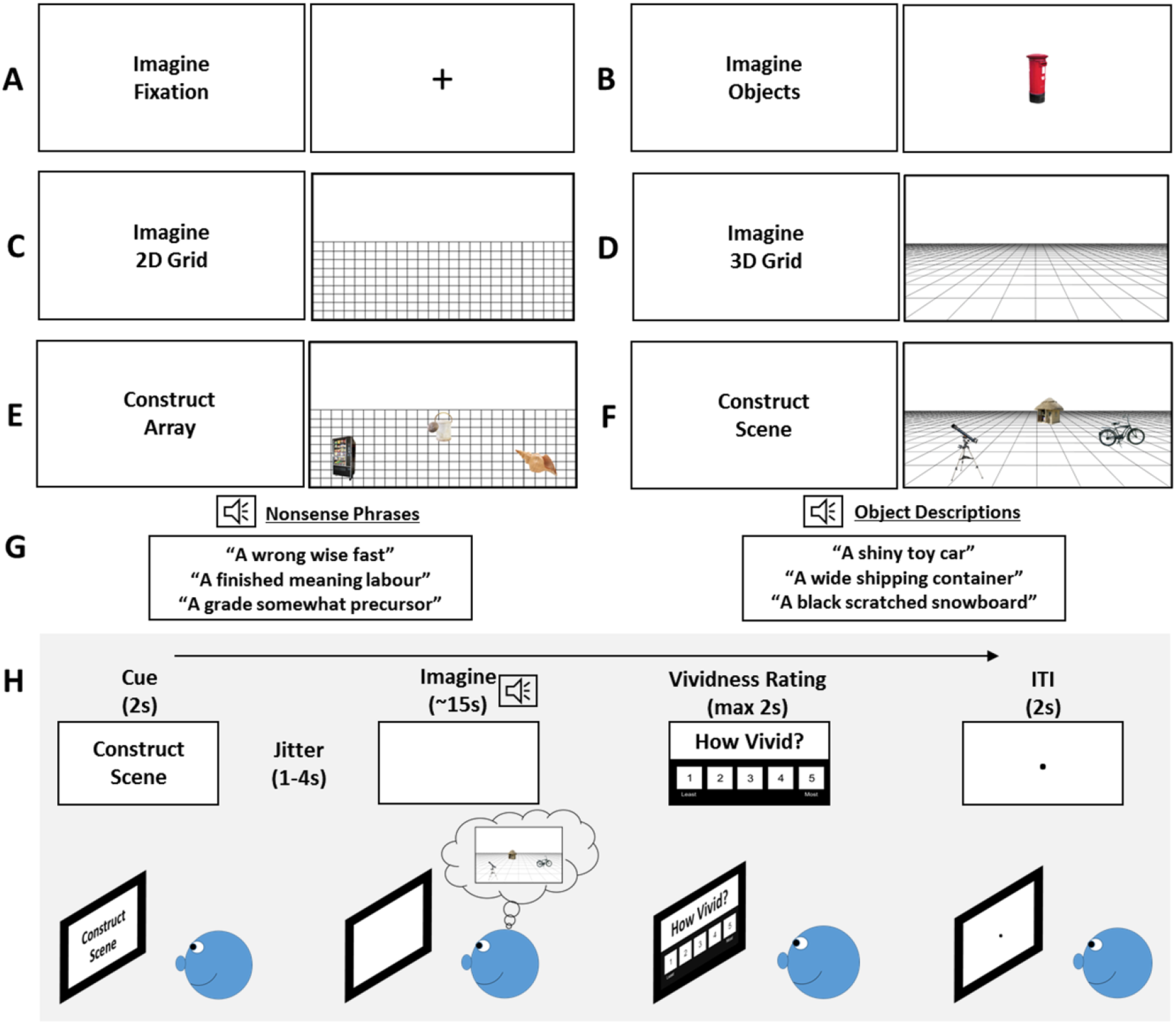
Experimental design. (A-F) Representations of how participants engaged in mental imagery during each task, with the text cues on the left of each panel. Note that participants looked at a blank white screen during the imagination phases. The images depicted on the right of each panel are based on what participants said they imagined during the task. (G) Examples of object descriptions and nonsense phrases. (H) Example of the time line of a single trial.

For the ‘Imagine Fixation’ task, participants were asked to imagine a black ‘plus’ sign in the centre of a blank white screen (Fig. 1A). While imagining the plus sign, participants were auditorily presented with three nonsense phrases (see Fig. 1G, left panel, for examples of the nonsense phrases), which compromised non-imageable abstract words, spoken one at a time. These were included in order to mirror the auditory input in the object tasks (see below) while precluding mental imagery. Participants were instructed not to try and interpret the nonsense phrases in any way. This ‘Imagine Fixation’ task was essentially a rest condition providing participants with a break from the more challenging imagination tasks described below.

For the ‘Imagine Objects’ task, participants were auditorily presented with three object descriptions (see Fig. 1G, right panel, for examples of object descriptions) one after another and instructed, when hearing the first object description, to imagine the object alone in the centre of the blank white screen (Fig. 1B). When hearing the second object description, they were asked to imagine the second object in place of the first in the centre of the screen and when hearing the third object description, to imagine it in place of the second object. During pre-scan training, participants were instructed and trained to imagine each object in complete isolation.

For the ‘Imagine 2D Grid’ task, participants were asked to create a mental image of a regular, flat 2D grid covering roughly the bottom two thirds of the blank screen (Fig. 1C). For the ‘Imagine 3D Grid’ task, participants were asked to create a mental image of a 3D grid covering roughly the bottom two thirds of the blank screen (see Fig. 1D). While imagining the grids, participants were auditorily presented with three nonsense phrases, spoken one at a time. The important difference between these tasks is that the 3D grid induces a sense of depth and 3D space.

For the ‘Construct Array’ task, participants were instructed to first imagine the 2D grid on the bottom two thirds of the screen. While doing this, participants were auditorily presented with three object descriptions one after another which they imagined on the 2D grid. More specifically, participants were asked, when hearing the first object description, to imagine the object in an arbitrary location on the 2D grid. When hearing the second object description, participants were asked to imagine it on another arbitrarily chosen location on the 2D grid while maintaining the image of the first object in its original location. When hearing the third object description, participants were asked to imagine it on another part of the 2D grid while maintaining the image of the first two objects in their original positions. We explicitly told participants that the final product of their mental imagery was to be three objects in random locations on a flat 2D grid (Fig. 1E).

For the ‘Construct Scene’ task, participants were instructed to first imagine a 3D grid on the bottom two thirds of the screen. While doing this, they were auditorily presented with three object descriptions one at a time which they were asked to imagine on the 3D grid. Specifically, participants were asked, when hearing the first object description, to imagine the object in any location on the 3D grid. When hearing the second object description, participants were asked to imagine it on another location on the 3D grid while maintaining the image of the first object in its original position. When hearing the third object description, participants were asked to imagine it on another part of the 3D grid while maintaining the image of the first two objects in their original locations. The final product of their mental imagery was to be 3 objects in a simple 3D scene (Fig. 1F).

For tasks which required object imagery (‘Imagine Objects’, ‘Construct Array’ and ‘Construct Scene’) we emphasised the importance of engaging imagination rather than memory for each object. We asked participants to imagine a unique version of each object based on the descriptions alone and, as far as possible, not to recall specific objects that they were familiar with, any personal memories involving the objects or any places that they might associate with the described objects. We also asked participants not to imagine any movement, even if objects had movable parts, but to create static images of each object in their mind’s eye.

For the Imagine 2D Grid and Imagine 3D Grid tasks, participants were instructed to keep their ‘viewpoint’ of the grid fixed and static and not to imagine either the grid moving or themselves moving along the grid. In contrast to the 2D grid, mental imagery of the 3D grid induces a sense of depth and participants were additionally asked not to imagine moving ‘into’ the 3D space in any way.

For the Construct Array and Construct Scene tasks, participants were asked that for each trial, they keep the objects separate from each other so that no objects physically touched and no objects interacted. We asked participants not to add any additional elements but to create the arrays and scenes using only the objects provided. Participants were asked to utilise the full extent of the grids to place the objects and to continue imagining the objects on the grids for the duration of the imagination period. Also, having imagined all three objects on the grid, participants were asked not to mentally ‘rearrange’ the objects. Rather, they were asked to leave them where they were initially placed in their mind’s eye. We asked participants to keep their viewpoint fixed and central and not to imagine themselves or any objects moving in any way. For the Construct Array task, we emphasised the importance of not linking the objects together into a scene but to arbitrarily place the objects in random locations.

Separate audio files were recorded for each object description and nonsense phrase. These were recorded in a sound proof room and spoken by a male voice in an even tone. Prior to the experiment, a separate group of participants (n = 10) rated each object description on whether it was space defining (SD) or space ambiguous (SA) (Mullally and Maguire, 2011, 2013) and also provided ratings of object permanence (Auger et al., 2012; Mullally and Maguire, 2011). Object descriptions and nonsense phrases were further rated for imageability. The auditory stimuli for each task were all three words in length and carefully matched on a range of specific features.

In relation to the object descriptions, the Imagine Objects, Construct Array and Construct Scene tasks were matched according to the number of SD and SA objects (F_(2,215)_ = 0.128, p = 0.880), ratings of object permanence (F_(2,215)_ = 0.106, p = 0.899), syllable number (F_(2,215)_ = 0.234, p = 0.792) and utterance length (F_(2,215)_ = 0.014, p = 0.986). In addition, the order of presentation of SD/SA items was balanced across all trials. Object triplets were arranged so that objects within each triplet had no obvious semantic or contextual associations.

In relation to nonsense phrases, syllable number (F_(2,215)_ = 1.953, p = 0.144) and utterance length (F_(2,215)_ = 0.591, p = 0.555) were matched across the Imagine Fixation, Imagine 2D Grid and Imagine 3D Grid tasks. In addition, syllable number (F_(5,431)_ = 0.925, p = 0.464) and utterance length (F_(5,431)_ = 0.658, p = 0.656) were matched across all tasks, and the nonsense phrases were rated as significantly less imageable than the object descriptions (t_(1,49)_ = 81.261, p < 0.001).

In summary, the two tasks of primary interest were the ‘Construct Array’ and ‘Construct Scene’ tasks. As described above, these tasks involved participants hearing three object descriptions and imagining them on either an imagined 2D or 3D space. With all else being equal in the stimuli, this simple manipulation of space gave rise to mental representations of non-scene arrays (objects and 2D space) and scenes (objects and 3D space). We also included tasks that examined the representation of three objects without a spatial context where the objects were imagined one after another in the same location on the centre of the screen, and the representation of either 2D or 3D space alone without objects. Overall, this novel paradigm allowed us to separately examine the neural correlates of constructing mental representations of objects alone (with no spatial context), two types of space (2D and 3D space alone with no objects) and two different combinations of objects and space where only one gave rise to scene representations. Importantly, no visual stimuli were presented during the imagination phase of any task (Fig. 1H). Therefore, between-task differences in neural recruitment could not be influenced by differences in visual input.

### Pre-scan training

Participants were trained before scanning to ensure task compliance. After listening to the instructions, participants engaged in four practice trials of each task while sitting at a desktop computer in a darkened room. They rated the vividness of mental imagery for each trial on a scale of 1 (not vivid at all)…5 (extremely vivid). If they gave a rating of 3 or below on any practice trial, they repeated the practice session. When participants rated 4 or above on all practice trials and indicated that they could vividly engage in the mental imagery relevant to each task, they were transferred to the scanner for the experiment.

### fMRI task

Each trial of the experiment (Fig. 1H) was comprised of a visual cue (2 seconds) which informed of the task, followed by a 1-4 second jitter and then the imagination phase of ∼15 seconds. During the imagination phase, participants engaged in the mental imagery pertinent to each task while hearing three auditory phrases (either objects or nonsense, depending on the task, Fig. 1G) delivered via MR compatible headphones (Sensimetrics model S14). The length of each auditory phrase was approximately 2s followed by a 1s gap between the presentation of each phrase. After hearing the third auditory phrase, participants had approximately 7s to finalise and maintain the mental image they had created. They were required to do this with their eyes open while looking at a blank white screen. They then rated the vividness of their mental image on a 1 (not vivid at all).5 (extremely vivid) scale (max 2 seconds). Finally, an inter-trial interval of 2 seconds preceded the cue for the next trial. Twenty four trials were presented for each condition (144 trials in total) in a pseudorandomised order across 4 separate blocks. Each block lasted approximately 15 minutes and blocks were separated by a brief rest period. It was emphasised to participants that the main objective of the experiment was to create vivid mental images in their imagination. However, to ensure participants were attending throughout, we included an additional 12 catch trials (2 per task) across the experiment where participants had to press a button if, within a nonsense or object triplet, they heard a repeated phrase.

### In-scan eye tracking

As a further measure of task compliance, we utilised in-scan eye tracking to ensure participants were performing each task according to the instructions. For the Imagine Fixation and Imagine Objects tasks, participants were asked to focus their eyes on the centre of the screen where they imagined the plus sign or objects to be. When imagining the 2D and 3D grids, they were asked to move their eyes around where they imagined the grids to be on the screen. For the Construct Array and Construct Scene tasks, participants were required to imagine each of three objects against the imagined 2D or 3D grid respectively. Eye tracking data were acquired using an Eyelink 1000 Plus (SR Research) eye tracker. The right eye was used for calibration, recording and analyses. During the imagination phase, the x and y coordinates of all fixations were recorded. Visualisation of fixation locations was performed with Eyelink Data Viewer (SR Research). Eye tracking data from 8 participants were unavailable due to technical issues, leaving 22 data sets in the eye-tracking analyses.

### Post-scan surprise memory tests

After completing the experiment and leaving the scanner, participants were taken to a testing room and given surprise item and associative yes/no recognition memory tests. Participants first performed an item memory test, where they were auditorily presented with all 216 object descriptions heard during the Imagine Object, Construct Array and Construct Scene tasks (72 objects per task) and an additional 72 lure items which were not heard during the experiment. Object descriptions were randomised across tasks and were presented one at a time. For each object description participants were asked to respond ‘yes’ if they thought the object description was heard during the scanning experiment and ‘no’ if they thought it was not.

Participants then performed a more difficult associative memory test. For this, participants were auditorily presented with 72 sets of object triplets (24 sets from each of the Imagine Object, Construct Array and Construct Scene tasks). Forty eight of these object triplets (16 from each of the three tasks) had been presented together during the experiment (intact triplets). Twenty four of the object triplets (8 from each of the three tasks) contained object descriptions that were presented during the experiment, but not together in a triplet (recombined triplets). For each object triplet, participants were asked to respond ‘yes’ if they thought the objects were heard together during the fMRI experiment and ‘no’ if they were not. For both memory tasks, participants also gave a confidence rating on a 1-5 scale for each decision and timing was self-paced (up to a maximum of 5 seconds each) for both the choices and confidence ratings. Note that we do not include the data from the confidence ratings from the associative memory test as they were, perhaps unsurprisingly, dominated by ‘guessing’ ratings. Memory test data from 7 participants were unavailable due to technical issues.

### Post-scan debriefing

Following the memory tests, participants were probed on how they approached each task, and performed a number of ratings as described in the Results section and Table 3.

**Table 3.** Subjective measures.

| **Question** | **Rating**<br>**(Yes/No or 1-5 Likert scale)** |  | **Tasks of**<br>**relevance** | **Mean ± SD** |
| --- | --- | --- | --- | --- |
| **How vivid was your imagination of the xxxx?**<br>(From the trial-by-trial vividness ratings during scanning) | 1 = not vivid at all | ...5 = extremely vivid | Imagine Fixation<br>Imagine Object<br>Imagine 2D grid<br>Imagine 3D grid<br>Construct Array<br>Construct Scene | 3.77 ± 0.65<br>4.16 ± 0.53<br>3.69 ± 0.62<br>3.84 ± 0.57<br>3.96 ± 0.52<br>4.03 ± 0.53 |
| **How difficult was it to imagine the xxxx?** | 1 = very easy | ...5 = very difficult | Imagine Fixation<br>Imagine Object<br>Imagine 2D grid<br>Imagine 3D grid<br>Construct Array<br>Construct Scene | 2.58 ± 1.22<br>2.01 ± 0.75<br>2.65 ± 1.09<br>2.13 ± 0.88<br>2.35 ± 0.76<br>2.43 ± 0.85 |
| **How often did you engage in off-task thoughts (mind-wander) during this task?** | 1 = never | ...5 = always | Imagine Fixation<br>Imagine Object<br>Imagine 2D grid<br>Imagine 3D grid<br>Construct Array<br>Construct Scene | 2.22 ± 0.80<br>1.77 ± 0.57<br>2.30 ± 0.99<br>1.87 ± 0.67<br>1.88 ± 0.65<br>1.95 ± 0.83 |
| **In general, how detailed do you think your images of the xxxx were?** | 1 = Hardly any details at all | ...5 = Extremely richly detailed | Imagine Object<br>Imagine 2D grid<br>Imagine 3D grid<br>Construct Array<br>Construct Scene | 3.27 ± 0.45<br>2.97 ± 0.81<br>3.22 ± 0.67<br>3.27 ± 0.69<br>3.45 ± 0.62 |
| **In general, did you feel that you successfully imagined the objects in a scene?** | 1 = never successful | . . .5 = always successful | Construct Scene | 3.77 ± 0.84 |
| **Did you feel that you imagined the objects in a scene at all and had to try to repress this?** | 1 = never had to repress | . . .5 = always repressed | Construct Array | 1.28 ± 0.69 |
| **On a scale of 1-5, how 2D/3D was your mental imagery during this task?** | 1 = completely 2D | . . .5 = completely 3D | Construct Array<br>Construct Scene | 1.90 ± 0.84<br>4.10 ± 0.82 |
| **Did you make up any stories or scenarios related to the xxxx?** | 0 = No<br>1 = Yes |  | Imagine Object | 0.02 ± 0.09 |
|  |  |  | Imagine 2D grid | 0.07 ± 0.25 |
|  |  |  | Imagine 3D grid | 0.02 ± 0.09 |
|  |  |  | Construct Array | 0.03 ± 0.18 |
|  |  |  | Construct Scene | 0.07 ± 0.25 |
| **Did you keep your viewpoint fixed?** | 0 = No<br>1 = Yes |  | Imagine 2D grid | 0.98 ± 0.09 |
|  |  |  | Imagine 3D grid | 0.97 ± 0.18 |
|  |  |  | Construct Array | 0.97 ± 0.18 |
|  |  |  | Construct Scene | 0.98 ± 0.09 |
| **To what extent did you imagine other objects or details in addition to the given objects? Did you. . . ?** | 1 = think of objects with other details the entire time | . . . 5 = not think of other objects or details at all | Imagine Object | 4.07 ± 0.83 |
|  |  |  | Construct Array | 4.23 ± 0.73 |
|  |  |  | Construct Scene | 4.08 ± 0.77 |

### Data acquisition and pre-processing

Structural and functional MRI data were acquired using a 3T Siemens Trio scanner (Siemens, Erlangen, Germany) with a 32-channel head coil within a partial volume centred on the temporal lobe and included the entire extent of the temporal lobe and all of our other regions of interest. Structural images were collected using a single-slab 3D T2-weighted turbo spin echo sequence with variable flip angles (SPACE) (Mugler et al., 2000) in combination with parallel imaging, to simultaneously achieve a high image resolution of ∼500 μm, high sampling efficiency and short scan time while maintaining a sufficient signal-to-noise ratio (SNR). After excitation of a single axial slab the image was read out with the following parameters: resolution = 0.52 × 0.52 × 0.5 mm^3^, matrix = 384 × 328, partitions = 104, partition thickness = 0.5 mm, partition oversampling = 15.4%, field of view = 200 × 171 mm 2, TE = 353 ms, TR = 3200 ms, GRAPPA x 2 in phase-encoding (PE) direction, bandwidth = 434 Hz/pixel, echo spacing = 4.98 ms, turbo factor in PE direction = 177, echo train duration = 881, averages = 1.9. For reduction of signal bias due to, for example, spatial variation in coil sensitivity profiles, the images were normalized using a prescan, and a weak intensity filter was applied as implemented by the scanner’s manufacturer. To improve the SNR of the anatomical image, three scans were acquired for each participant, coregistered and averaged. Additionally, a whole brain 3D FLASH structural scan was acquired with a resolution of 1 × 1 × 1 mm.

Functional data were acquired using a 3D echo planar imaging (EPI) sequence which has been demonstrated to yield improved BOLD sensitivity compared to 2D EPI acquisitions (Lutti et al., 2013). Image resolution was 1.5mm^3^ isotropic and the field-of-view was 192mm in-plane. Forty slices were acquired with 20% oversampling to avoid wrap-around artefacts due to imperfect slab excitation profile. The echo time (TE) was 37.30 ms and the volume repetition time (TR) was 3.65s. Parallel imaging with GRAPPA image reconstruction (Griswold et al., 2002) acceleration factor 2 along the phase-encoding direction was used to minimise image distortions and yield optimal BOLD sensitivity. The dummy volumes necessary to reach steady state and the GRAPPA reconstruction kernel were acquired prior to the acquisition of the image data as described in Lutti et al. (2013). Correction of the distortions in the EPI images was implemented using B0-field maps obtained from double-echo FLASH acquisitions (matrix size 64×64; 64 slices; spatial resolution 3mm^3^; short TE=10 ms; long TE=12.46 ms; TR=1020 ms) and processed using the FieldMap toolbox in SPM (Hutton et al., 2002).

Preprocessing of structural and fMRI data was conducted using SPM12 (Wellcome Centre for Human Neuroimaging, University College London). All images were first bias corrected, to compensate for image inhomogeneity associated with the 32 channel head coil (Van Leemput et al., 1999). Fieldmaps were collected and used to generate voxel displacement maps. EPIs for each session were then realigned to the first image and unwarped using the voxel displacement maps calculated above. The three high-resolution structural images were averaged to reduce noise, and co-registered to the whole brain structural FLASH scan. EPIs were also co-registered to the whole brain structural scan and spatially smoothed using a Gaussian smoothing kernel of 4 × 4 × 4 mm full-width at half maximum.

### Statistical analyses: behavioural data

Data from eye tracking, in-scan vividness ratings, post-scan memory tests and debrief ratings were analysed using repeated measures ANOVAs (SPSS 17.0, Chicago: SPSS inc.) with a significance threshold of p < 0.05. Where Mauchly’s test indicated that the assumption of sphericity had been violated, degrees of freedom were corrected using Greenhouse-Geisser estimates of sphericity.

### Statistical analyses: fMRI data

We used non-rotated task Partial Least Squares (PLS) for data analysis which is a multivariate method for extracting distributed signal changes related to varying task demands (Krishnan et al., 2011; McIntosh and Lobaugh, 2004). Data for each condition were included in a block design analysis and we conducted separate analyses for each of our contrasts of interest. Significance for each contrast was independently determined using a permutation test with 1000 permutations. We considered latent variables less than p = 0.05 as significant. The reliability of voxel saliences was assessed by means of a bootstrap estimation of the standard error (McIntosh and Lobaugh, 2004). Bootstrapping is a sampling technique in which subjects are randomly selected into the analysis with replacement from the entire group of subjects. For each new sample, the entire PLS analysis is recalculated. In the present study, this sampling and analysis procedure was carried out 500 times, resulting in estimates of the standard error of the salience at each voxel. No corrections for multiple comparisons are necessary because the voxel saliences are calculated in a single mathematical step on the whole volume. We considered clusters of 10 or more voxels in which the bootstrap ratio was greater than 1.96 (approximately equivalent to p < 0.05) to represent reliable voxels. In the current analyses, we specified a 14.6 second temporal window for each event (i.e. 4 TRs) to include the active phase of mental construction. Importantly, for each significant contrast reported in the main text, confidence intervals did not cross the 0 line suggesting that each condition contributed to the pattern.

We used a large region of interest that included the whole medial temporal lobe (MTL) – hippocampus, entorhinal cortex (ENT), PRC, as well as PHC, RSC and PCC which have been implicated in scene processing. The mask also extended posteriorly to encompass regions of the early visual cortices (EVC) including the precuneus (only inferior portions due to the partial volume), the calcarine sulcus, the lingual gyrus and portions of the posterior fusiform cortex, given these regions have previously been implicated in different elements of mental imagery (de Gelder et al., 2016; Lambert et al., 2002; Klein et al., 2000).

## Results

Our main focus was on contrasts involving the Construct Scene task (the full results of all the task comparisons are provided in Table 1). The contrast of primary interest was the Construct Scene task with the closely matched Construct Array task. As described above, these two conditions were well matched, requiring mental construction and associative processing of objects and space. The only difference between them was the scene construction task required objects to be imagined on a 3D grid that gave rise to a scene-like representation (compare the panels in Fig. 1E and F). Directly contrasting these tasks, therefore, allowed us to investigate brain regions that underpin scene construction while controlling for content, mental constructive and associative processes.

**Table 1.**
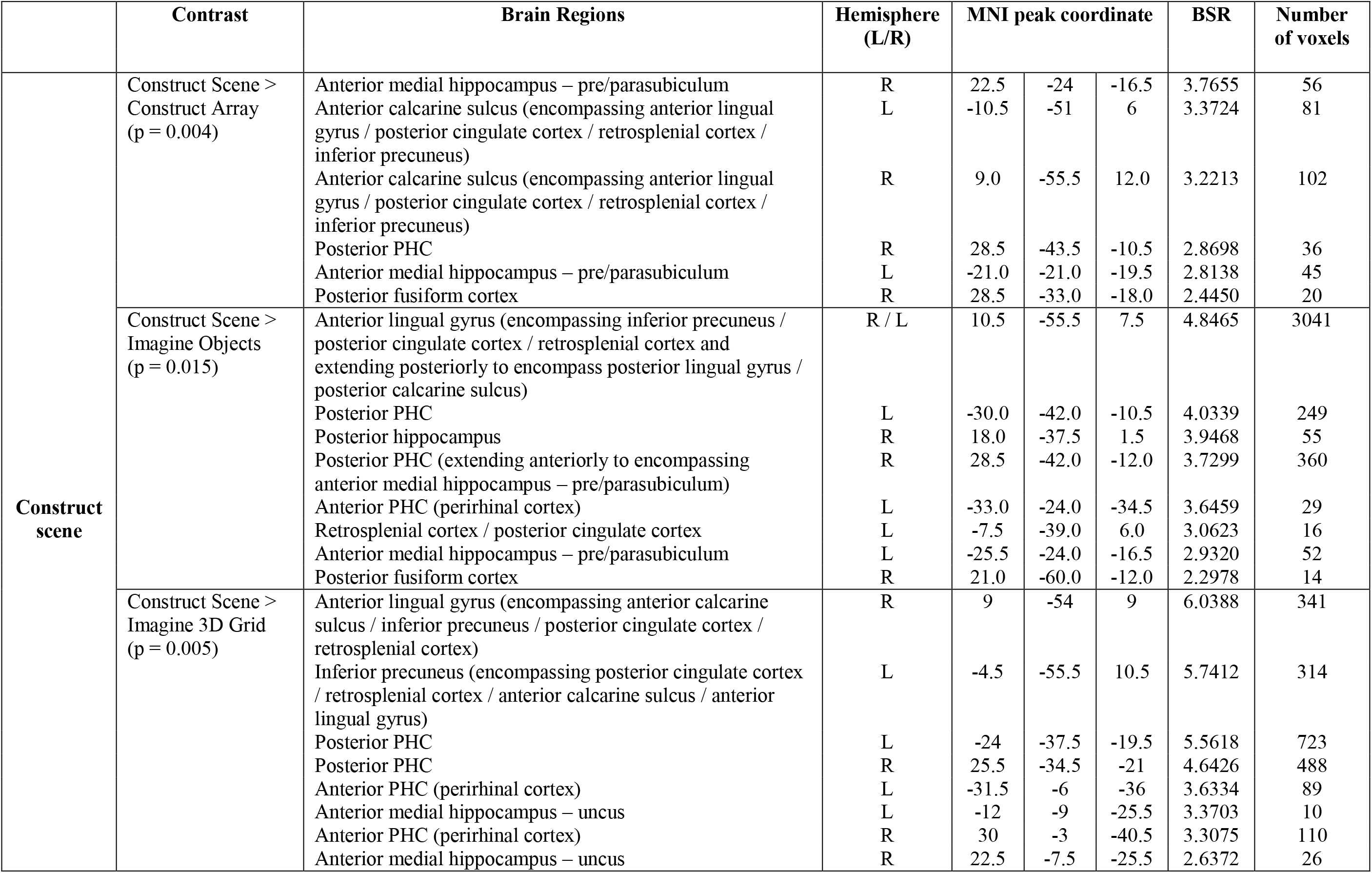
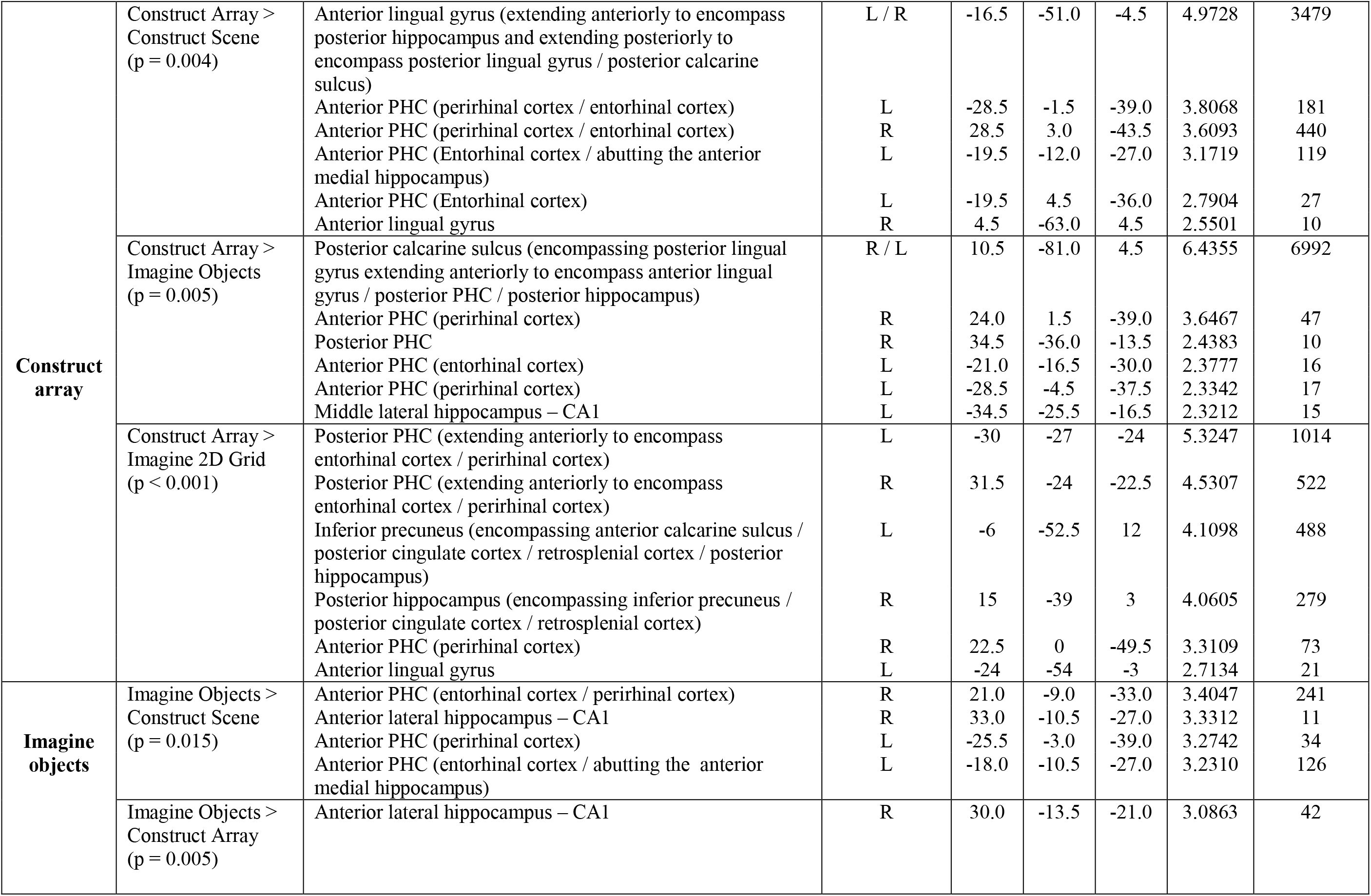
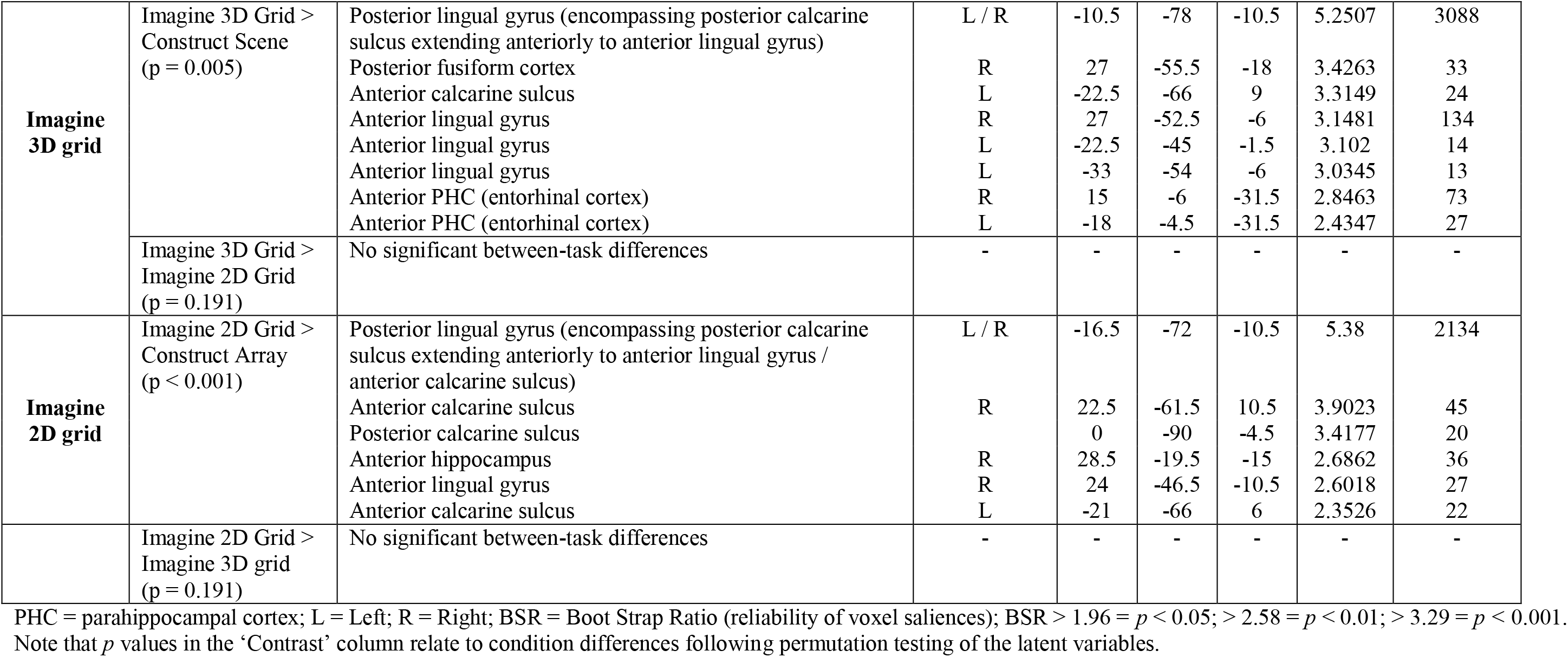
Results of the fMRI task comparisons.

### fMRI task comparisons

Comparison of the Construct Scene task with the Construct Array task revealed that, in line with our prediction, a circumscribed region of the anterior medial hippocampus (peak voxel at y = −24) encompassing the pre/parasubiculum was preferentially recruited, bilaterally, during scene construction along with the PHC, RSC and PCC (Fig. 2B, Fig. 3). Interestingly, the opposite contrast showed that array, more than scene, construction engaged bilateral ENT, PRC, EVC and the left posterior hippocampus which was part of a larger cluster of activity which encompassed the anterior lingual gyrus and portions of the EVC. This contrast also revealed activation of the left ENT abutting the anterior medial hippocampus (peak voxel at y = −12) that was more anterior to the pre/parasubiculum engaged by scene construction (Fig. 2B, Fig. 3).

**Figure 2.**
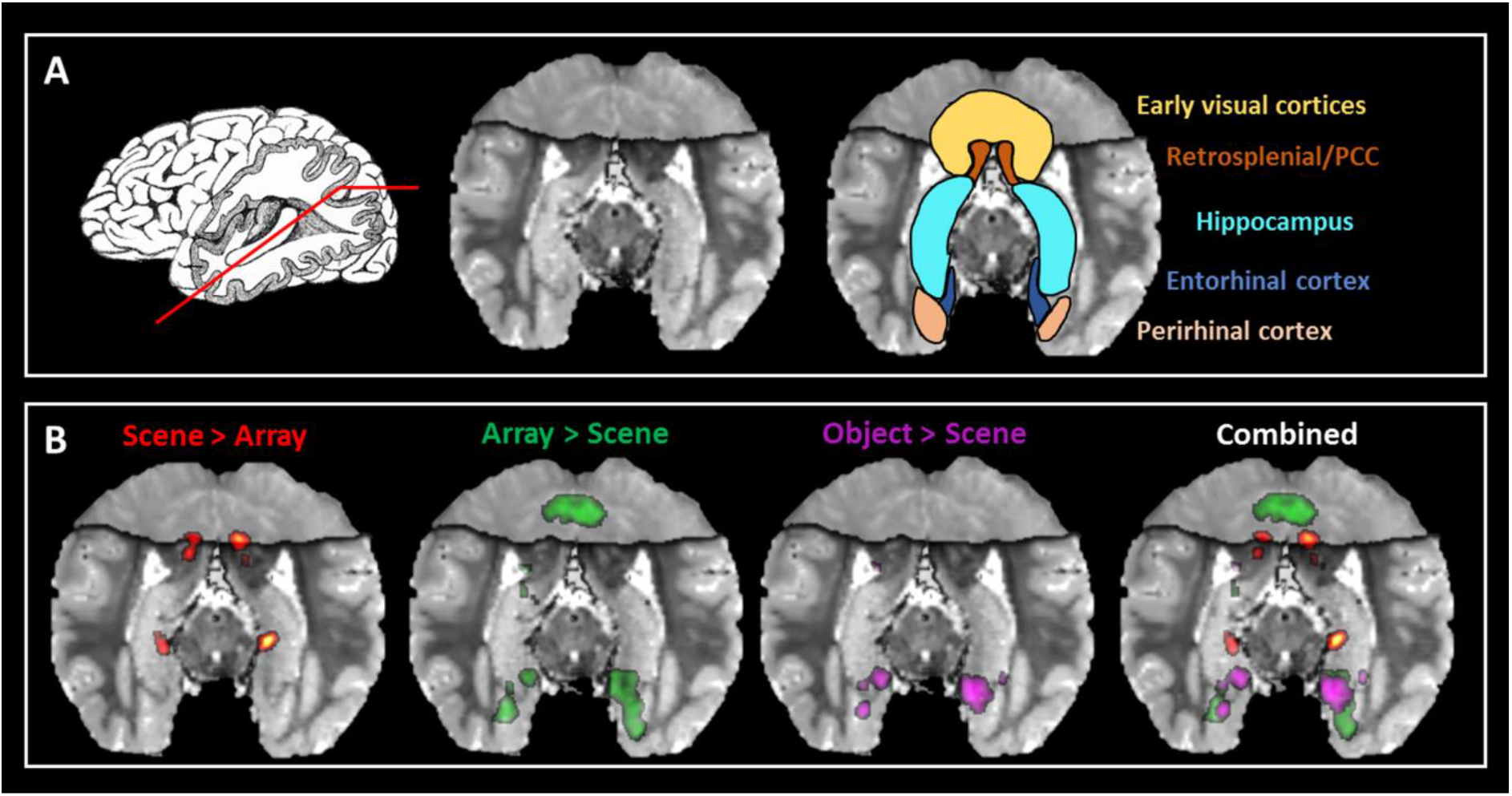
Results of the fMRI analysis. (A) Left panel, a representation of the oblique angle cutting through the hippocampus that we use to present the results in the other panels. Middle panel, shows the averaged structural MR image of the participants exposing the length of the hippocampus. Right panel, depicts the regions of particular interest. (B) From left to right, the results for the contrast of Construct Scene > Construct Array, Construct Array > Construct Scene, Imagine Object > Construct Scene and all of the results combined. PCC = posterior cingulate cortex; results are thresholded at p < 0.05.

**Figure 3.**
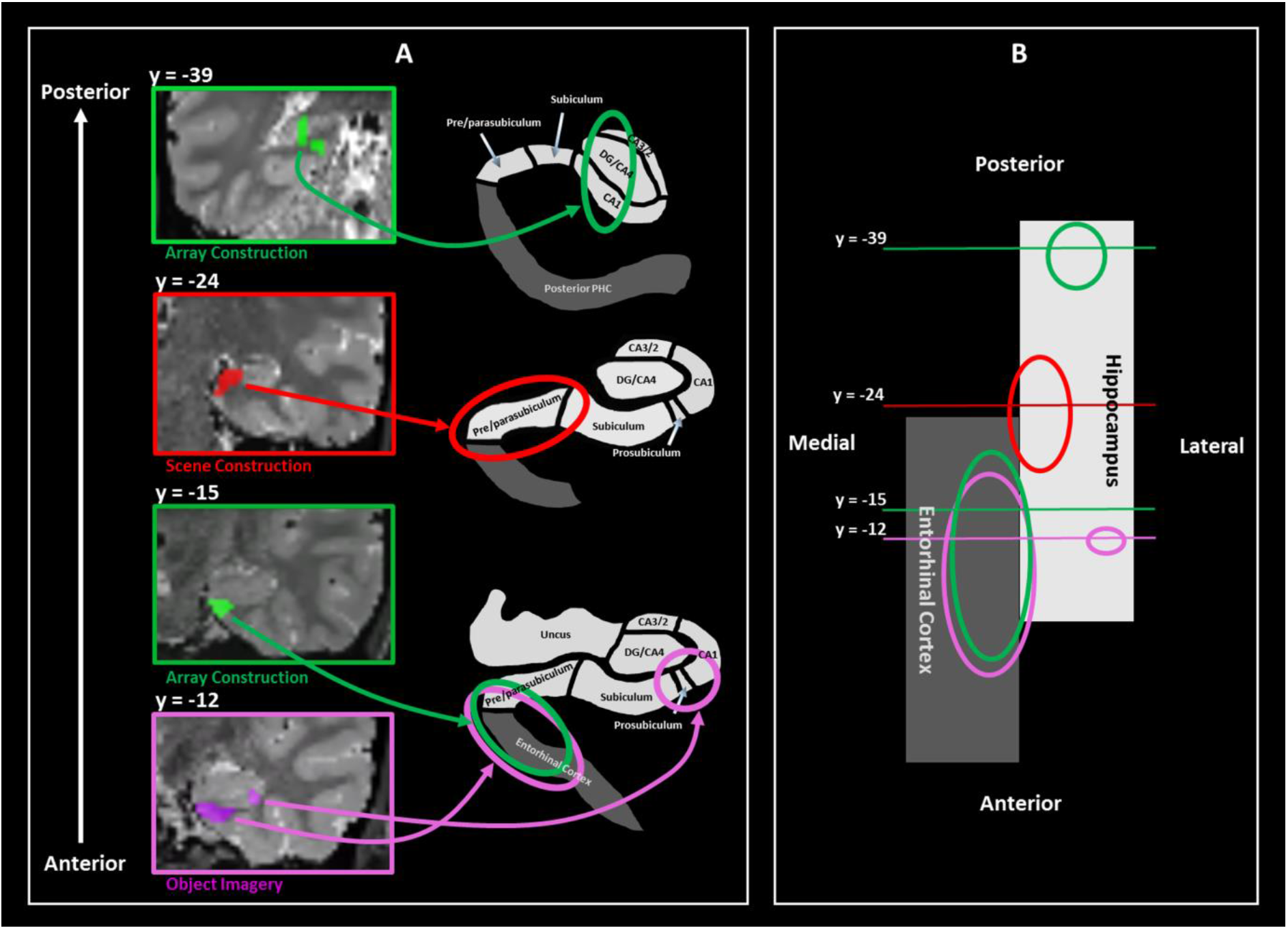
Summary of the main hippocampal activations. (A) The location of the left posterior hippocampal activation for the Construct Array > Construct Scene contrast (left, top panel). The right anterior medial hippocampal activation – pre/parasubiculum – observed for the contrast of Construct Scene > Construct Array (left, second panel from top). The entorhinal region abutting the much more anterior pre/parasubiculum that was recruited for both Construct Array (left, third panel from top) and Imagine Objects (left, bottom panel) tasks more so than Construct Scenes. The right anterior lateral hippocampal activation – prosubiculum/CA1 – for the contrast of Imagine Objects > Construct Scene (left, bottom panel). The right panels show representative schematics of the locations of the hippocampal subregions. (B) Schematic representation of the hippocampus (white) and entorhinal cortex (grey) in the axial plane. The location of each of the coronal plane images presented in (A) is shown along with representations of the extent of each cluster.

The contrast of Construct Scene with Imagine Objects provided further support that bilateral pre/parasubiculum along with the PHC, RSC and PCC were specifically associated with scene construction (Fig. 3, Table 1,). The reverse contrast showed that the mental construction of objects, more so than scenes, was associated with bilateral PRC and ENT. The right anterior lateral hippocampus, encompassing prosubiculum/CA1, and a left ENT activation that abutted the anterior medial hippocampus were also engaged. This area was more anterior (peak voxel at y = −10.5) to that associated with scene construction (Fig. 2B, Fig. 3).

The contrast of Construct Scenes with the Imagine 3D Grid revealed increased engagement of an anterior medial portion of the hippocampus in the approximate location of the uncus (peak voxel at y = −9) and bilateral PRC. The reverse contrast showed that the mental construction of 3D grids, more so than scenes, was associated with bilateral ENT and posterior portions of EVC. Imagine 3D Grid did not evoke increased hippocampal activation compared to Imagine 2D Grid, suggesting that 3D space alone was insufficient to engage the hippocampus.

To summarise (see Fig. 3), our experimental design allowed us to parse the hippocampus and related areas dependent on the process that was being engaged. Circumscribed portions of the bilateral pre/parasubiculum (around y = −24) were specifically recruited during scene construction. By contrast, a more anterior portion of the ENT that abutted the anterior medial hippocampus was engaged during both array and object construction. Of note, these activated regions were clearly distinct (explicit smoothing = 4mm; Euclidean distance between peak voxels of the Construct Scene versus Construct Array contrasts 11.89mm; Construct Array versus Imagine Object contrasts 13.64mm). The construction of mental images of arrays was also associated with increased activity in the posterior hippocampus as part of a larger cluster which encompassed the lingual gyrus and EVC. Object construction engaged anterior lateral hippocampus in the region of prosubiculum/CA1. Outside of the hippocampus, compared to array construction, the PHC, RSC and PCC were preferentially recruited during scene construction. In contrast, compared to scene construction, array construction was associated with more posterior portions of the EVC, while both array and object construction were more strongly associated with the ENT and PRC.

But could other factors have influenced the results? We conducted a range of further analyses to investigate.

### Did participants truly engage with the tasks?

The construction of mental imagery cannot be directly measured. We therefore used a combination of methods to assess task attentiveness and compliance. First, we included catch trials where participants had to press a button if, during any trial, they heard a repeated phrase. On average, 94% (SD = 0.09) of catch trials were correctly identified indicating that participants remained attentive throughout the experiment.

Second, we utilised in-scan eye tracking to ensure participants were performing each task according to the instructions. Visualisation of fixations confirmed that participants engaged in each task according to our instructions (see examples in Fig. 4A). To formally determine the extent of eye movements, we measured the variance of all fixations in the horizontal axis during the construction phase of each trial (Fig. 4B). We predicted that if participants performed the mental imagery tasks as expected, there would be less variance in fixation location during the centre-focussed Imagine Fixation and Imagine Objects tasks and a more dispersed pattern of fixations across the other tasks. The results of a repeated measures ANOVA revealed a significant difference in the variance of fixations between tasks (F_(2.28,47.77)_ = 26.22, p < 0.001). In line with our prediction, post hoc analyses revealed significantly less variance during the Imagine Fixation task compared to the other tasks (compared to the Imagine 2D Grid t_(21)_ = 6.286, p < 0.001; Imagine 3D Grid t_(21)_ = 5.296, p < 0.001; Construct Array t_(21)_ = 6.247, p = 0.001; Construct Scene t_(21)_ = 5.839, p < 0.001). Significantly less variance was also observed in the Imagine Objects task compared to the other tasks (Imagine 2D Grid t_(21)_ = 6.241, p < 0.001; Imagine 3D Grid t_(21)_ = 4.949, p < 0.001; Construct Array t_(21)_ = 6.266, p < 0.001; Construct Scene t_(21)_ = 5.336, p < 0.001). There was no difference between the Imagine Fixation and Imagine Objects tasks (t_(21)_ = 1.702, p = 0.806). Variance during the Imagine 2D Grid task was significantly less than during the Imagine 3D Grid task (t_(21)_ = 3.819, p = 0.015). No other significant between-task differences were observed, including between the Construct Scene and Construct Array tasks (t_(21)_ = 1.897, p = 0.884).

**Figure 4.**
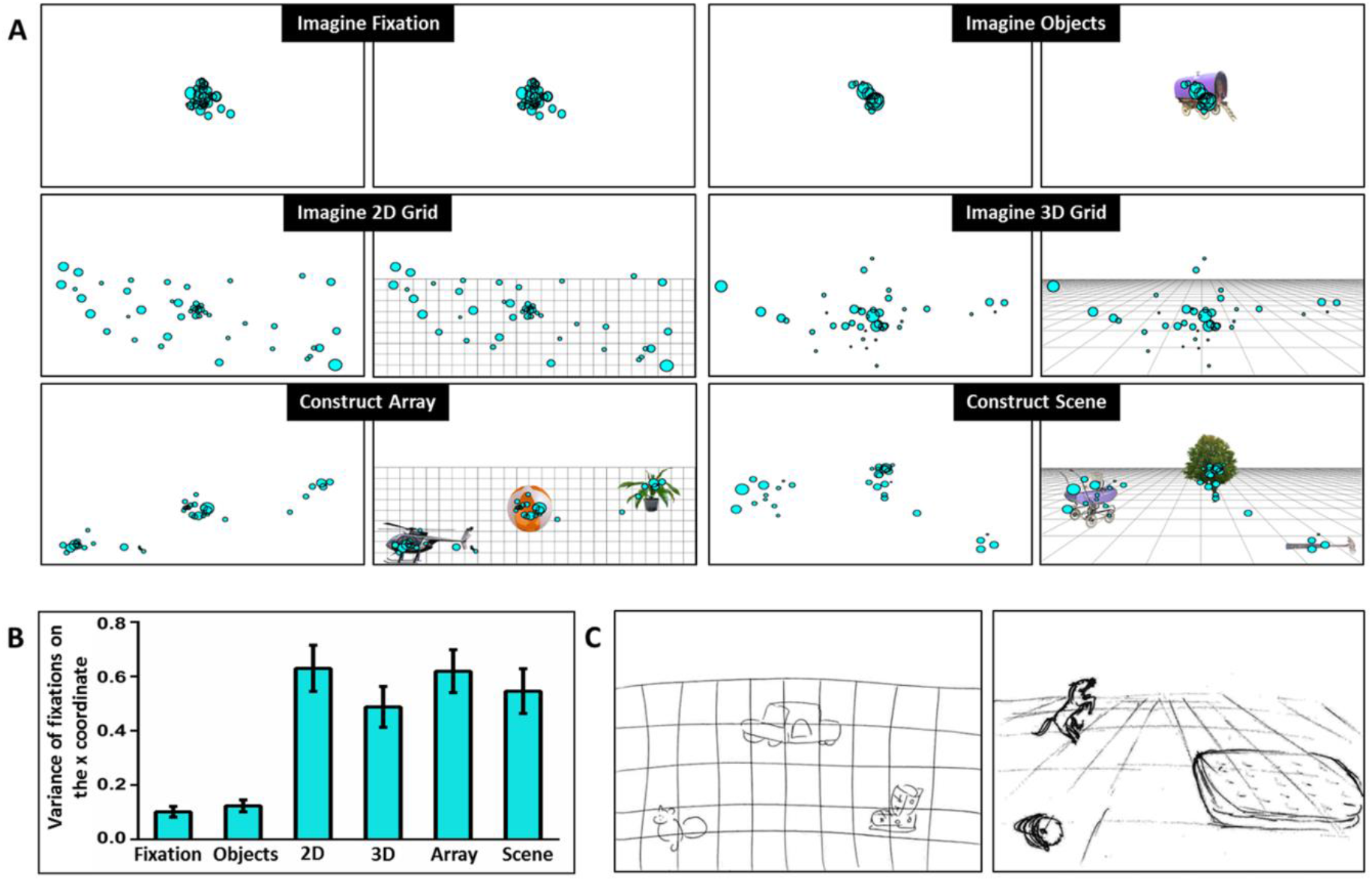
Eye movements and examples of post-scan drawings. (A) Representative examples of fixation locations (cyan circles) during the imagination phase of a single trial of each task are overlaid on a blank white screen on which participants focused their imagination (left panel). A visual representation of what participants were asked to imagine on the screen is shown to the right of each panel. Note the central focus of fixations for the Imagine Fixation and Imagine Objects tasks, the more dispersed pattern over the bottom two thirds of the screen for the Imagine 2D Grid and Imagine 3D Grid tasks and the three clusters for both the Construct Array and Construct Scene tasks representing the locations of the imagined objects. (B) The mean variance (+/− 1 SEM) of fixations on the x coordinate during the imagination phase of each task is shown. (C) Representative examples of post-scan drawings for the Construct Array (left) and Construct Scene (right) tasks.

Taken together, these measures provide quantitative evidence that participants paid attention during the experiment and engaged in mental imagery construction in accordance with task instructions.

After scanning, we also asked participants to draw how they had imagined the arrays and scenes during the fMRI tasks. Examples are shown in Fig. 4C and also indicate that participants complied with task requirements. The drawings also show that, despite both being composed of three objects related to a space, there was a clear representational difference between the arrays and the scenes. Further informative measures were obtained during post-scan testing and debriefing, and these are described in following sections.

### Did other eye movement features contribute to between-task differences?

To investigate the possibility that between-task differences in neural recruitment may be explained by other eye movement features, we investigated the number of fixations, fixation durations, the number of saccades, saccade amplitudes and scan paths, with a specific focus on our tasks of interest, Construct Array and Construct Scene. There were no differences in terms of the number of fixations (t_(21)_ = 0.144, p = 1.00), fixation durations (t_(21)_ = 0.423, p = 1.00), the number of saccades (t_(21)_ = 0.033, p = 1.00) or saccade amplitudes (t_(21)_ = 1.822, p = 0.726).

To examine scan paths, we split the screen into three equal areas of interest (AOI), left, middle and right, and plotted the scan path for each trial. Two variables were measured in order to provide an index of the spatial distribution of scanning - the number of fixations and the dwell time within each AOI. Analyses showed that there was a task by AOI interaction for number of fixations (F_(1,249,26.237)_ = 5.989, p = 0.016), with Construct Array associated with more fixations to the right of the screen (t_(21)_ = 2.251, p = 0.035, d = 0.16), Construct Scenes with more fixations in the middle (t_(21)_ = 2.175, p = 0.041, d = 0.24), with no difference for the left side of the screen (t_(21)_ = 1.784, p = 0.089). Neither of these effects survived Bonferroni correction, and the effect sizes (Cohen’s d) were small. There was also a task by AOI interaction for dwell time (F_(1.329,27.918)_ = 4.0, p = 0.045), with Construct Scene associated with a longer dwell time in the middle (t_(21)_ = 2.369, p = 0.027, d = 0.19), with no difference for the left of the screen (t_(21)_ = 0.772, p = 0.449) or the right of the screen (t_(21)_ = 1.840, p = 0.080). This effect did not survive Bonferroni correction, and the effect size was small.

Overall, therefore, the Construct Array and Construct Scene tasks were generally well matched, making it unlikely that between-task differences in neural recruitment were related to eye movements.

### Was hippocampal recruitment related to mnemonic processing?

Once out of the scanner after completing the experiment, participants were given two surprise memory tests (see Materials and Methods). Given the large number of stimuli, and the fact that there was no explicit instruction to encode information – rather the emphasis was on mental construction, and memory was never mentioned – our expectation was that performance would be poor, even at chance, on the memory tests. We felt it was necessary, however, to conduct these tests in case successful encoding differed across tasks, and this could have explained differences in the brain areas that were engaged. Scores (Means, SD) are shown in Table 2.

**Table 2.** Item and associative memory test performance (% correct).

|  | **Task** | **Mean ± standard deviation** |
| --- | --- | --- |
| **Item memory** | Imagine Object | 82.85 ± 11.68 |
| | Construct Array | 76.03 ± 13.47 |
| | Construct Scene | 71.07 ± 11.87 |
| | Identifying Novel Items | 81.94 ± 11.17 |
| **Associative memory intact triplets** | Imagine Object | 51.09 ± 14.18 |
| | Construct Array | 55.16 ± 21.04 |
| | Construct Scene | 51.90 ± 13.38 |
| **Associative memory recombined triplets** | Imagine Object | 46.20 ± 17.45 |
| | Construct Array | 54.35 ± 19.44 |
| | Construct Scene | 48.91 ± 19.91 |

For the item memory test, participants performed above chance at recalling stimuli that were from the Imagine Objects (t_(22)_ = 13.491, p < 0.001), Construct Array (t_(22)_ = 9.268, p < 0.001) and Construct Scene (t_(22)_ = 8.514, p < 0.001) tasks, and were above chance at identifying novel items (t_(22)_ = 13.710, p < 0.001). The good performance on this test (with task means between 70-83% correct – see Table 2) is a further indication that the participants paid attention, encoded information and were engaged by the tasks. A repeated measures ANOVA revealed a significant between-task effect on the item memory test (F_(1.77, 38.98)_ = 9.524, p < 0.001). Post hoc analyses showed that participants were better at recognising object descriptions which were presented in the Imagine Objects task than objects presented in the Construct Array (t_(22)_ = 4.829, p < 0.001) and Construct Scene (t_(22)_ = 7.210, p < 0.001) tasks. Participants were also better at recognising novel items than objects presented in the Construct Scene task (t_(22)_ = −3.382, p = 0.016). Notably, there was no significant difference between the Construct Array and Construct Scene tasks (t_(22)_ = 2.707, p = 0.075).

On the very challenging associative memory task, as we expected, participants did not perform above chance on the recognition of intact triplets that were from the Imagine Objects (t_(22)_ = 0.368, p = 0.717), Construct Array (t_(22)_ = 1.177, p = 0.252) and Construct Scene (t_(22)_ = 0.682, p = 0.503) tasks. A repeated measures ANOVA showed that there were no significant differences between the tasks for either the recognition of intact object triplets (F_(2,44)_ = 0.870, p = 0.426), or correct rejection of recombined object triplets (F_(2,44)_ = 1.651, p = 0.204). This included no significant difference between Construct Array and Construct Scene tasks (intact triplets: t_(22)_ = 1.047, p = 0.667; recombined triplets: t_(22)_ = 1.124, p = 0.616).

Overall, these results revealed no significant differences in memory performance in particular between our two tasks of interest, Construct Array and Construct Scene. Therefore, the differences we observed in neural recruitment during fMRI cannot be accounted for by differences in encoding success. Of note, it was not appropriate to run a subsequent memory analysis on the fMRI data using the individual object stimuli. This is because the three object descriptions that comprised one trial were presented in quick succession and it was not possible with fMRI to reliably tease apart signals relating to the specific items within a trial. Considering associative memory for the triplets, given that performance was not above chance in the subsequent surprise memory test, and that participants expressed low confidence about their memory decisions, using these data to interpret the fMRI data would be ill-advised. Moreover, in the associative memory test, two thirds of the triplets were tested intact, but one third of triplets were recombined in order to act as lures. Therefore, any subsequent memory fMRI analysis would likely be underpowered.

### Can imagery vividness account for hippocampal engagement?

During fMRI scanning, participants rated the vividness of mental imagery on each trial (see Materials and Methods; Fig. 1H; means (SDs) on Table 3). Results of the repeated measures ANOVA revealed a significant between-tasks difference in vividness ratings (F_(2.70, 78.26)_ = 11.60, p < 0.001). Post hoc analyses showed that mental imagery during the Imagine Objects task was rated as more vivid than during the Imagine Fixation (t_(29)_ = 4.095, p = 0.005), Imagine 2D Grid (t_(29)_ = 5.586, p < 0.001), Imagine 3D Grid (t_(29)_ = 4.195, p = 0.004), Construct Array (t_(29)_ = 4.506, p < 0.001) and Construct Scene (t_(29)_ = 3.265, p = 0.041) tasks. Imagery during the Construct Array task was rated significantly more vivid than during the Imagine 2D Grid task (t_(29)_ = 3.398, p = 0.029). Imagery during the Construct Scene task was rated significantly more vivid than during the Imagine 2D Grid (t_(29)_ = 4.116, p = 0.004) and Imagine 3D Grid (t_(29)_ = 3.224, p = 0.046) tasks. Importantly, no significant difference was observed between the Construct Array and Construct Scene tasks (t_(29)_ 2.116, p = 0.483).

### Can perceived task difficulty or subjective differences in mental imagery account for hippocampal recruitment?

In the debriefing session after scanning, and once the surprise memory tests were completed, participants were asked about their experience of performing each task (see means (SDs) on Table 3). Participants reported that they could perform the tasks with ease with no between-task differences for perceived task difficulty (F_(3.37,97.85)_ = 2.396, p = 0.066; including no difference between the Construct Array and Construct Scene tasks, t(29) = 0.524, p = 1.00). Significant between-task differences were observed for the rating of mind-wandering (F(3.46,100.39) = 3.638, p = 0.011). Post hoc analyses showed that compared to the Imagine Objects task, participants were more prone to engage in mind-wandering during the Imagine Fixation (t(29) = 3.465, p = 0.025) task. This makes sense considering the fixation task was included as a rest condition for participants. There was no significant difference between Construct Array and Construct Scene tasks (t(29) = 0.436, p = 1.00). Significant differences were also observed on the rating of detail of mental imagery (F_(3.47, 100.70)_ = 3.510, p = 0.014). Post hoc analyses showed that mental imagery during the Construct Scene task was significantly more detailed than during the ‘Imagine 2D Grid’ task (t_(29)_ = 3.093, p = 0.043). No other significant between-task differences were observed, including between Construct Array and Construct Scene tasks (t_(29)_ = 1.884, p = 0.514).

Participants further confirmed (Table 3) that, during the Construct Scene task, they were successful at creating a scene in their minds eye. In contrast, participants reported a clear sense of imagining objects on a 2D grid during the Construct Array task and stated that they rarely felt a need to repress mental imagery of scenes during this task. Direct comparison showed that, as expected, the Construct Scene task was rated as subjectively more 3D than the Construct Array task which was rated as more 2D (t_(29)_ = 11.988, p < 0.001). There were no significant differences between the tasks (including between Construct Array and Construct Scenes tasks) on several other subjective measures: the creation of narratives about the stimuli (F(2.15,62.43) = 0.597, p = 0.565; Construct Array versus Construct Scene t_(29)_ = 1.00, p = 0.981), the fixedness of the viewpoint (F_(1.96,56.75)_ = 0.139, p = 0.867; Construct Array versus Construct Scene t(29) = 0.441, p = 0.999) and the inclusion of extraneous objects or other details (F_(2,58)_ = 0.957, p = 0.390; Construct Array versus Construct Scene t(29) = 1.055, p = 0.657).

In summary, subjective measures indicated that the participants performed the task with ease, and adhered to the instructions. As might be expected, there were some minor differences, for example increased mind-wandering during the Imagine Fixation task but, importantly, no significant differences between the Construct Array and Construct Scene tasks.

### Results summary

The results of the fMRI analyses revealed that when other associative and mental constructive processes were taken into account, a specific region of the anterior medial hippocampus – corresponding to the location of the pre/parasubiculum – was engaged during scene construction along with other regions which have previously been linked to scene processing including the PHC, RSC and PCC. In contrast, array construction was more strongly associated with the ENT, PRC, posterior portions of EVC and activation within the left posterior hippocampus and left ENT abutting the anterior medial hippocampus. Importantly, this latter activation was in a location more anterior to the cluster observed during scene construction. Interestingly, the Imagine Objects task resulted in activation of the anterior lateral hippocampus. The differing patterns of neural recruitment between the very tightly matched Construct Array and Construct Scene tasks could not be accounted for by differences in eye movements, mnemonic processing or the phenomenology of mental imagery.

## Discussion

The aim of this study was to compare accounts that place associative processes at the heart of hippocampal function with the theory that proposes scene construction as one of its primary roles. Using closely matched tasks during high resolution fMRI we found that, as predicted, the pre/parasubiculum in the anterior medial hippocampus was preferentially engaged by the construction of scenes (three objects and a 3D space). However, it was also evident that different regions *within* the hippocampus were engaged by the construction of arrays (three objects and a 2D space) that did not evoke scene representations. In this case, the posterior hippocampus and an ENT region that abutted the anterior hippocampus were recruited. Even the imagination of object triplets that had no spatial context activated this latter region along with an anterior portion of the lateral hippocampus in the approximate location of prosubiculum/CA1. Overall, these results show that one possible reason for ongoing debates about how the hippocampus operates may be because it does not only process space or associations or scenes. Instead, there may be multiple processing circuits within the hippocampus that become engaged depending on task demands.

Our primary contrast of interest was array construction compared with scene construction. These tasks were closely matched on stimulus content and mental constructive and associative processes. Attention, eye movements, encoding success and perceived difficulty did not differ between them. Phenomenologically, the vividness and detail of their imagery were also matched. Nevertheless, in line with our prediction and previous reports, a circumscribed portion of the pre/parasubiculum in the anterior medial hippocampus was preferentially involved in scene construction (Zeidman and Maguire, 2016; Zeidman et al., 2015a; Zeidman et al., 2015b; Hodgetts et al., 2016; Maas et al., 2014). Importantly, our findings reveal for the first time that this region is preferentially recruited, not for mental construction per se, not for imagining a 3D space alone, but specifically for mental construction of scenes that are couched within a naturalistic 3D spatial framework.

Drawing on the latest anatomical evidence, Dalton and Maguire (2017) recently noted that the pre/parasubiculum is a primary target of the parieto-medial temporal processing pathway and may receive integrated information from foveal and peripheral visual inputs (Kravitz et al., 2011). Thus, it has privileged access to holistic representations of the environment and so could be neuroanatomically determined to preferentially process scenes. Indeed, Dalton and Maguire (2017) suggest the pre/parasubiculum may be the hippocampal ‘hub’ for scene-based cognition. The PHC, RSC and PCC are also implicated in the anatomical scene processing network connecting with the pre/parasubiculum. Aligning with this evidence and their known links with scene processing (Epstein, 2008), we found that these brain areas were more engaged during scene compared to array construction.

By contrast, array construction engaged a different set of brain areas, namely, the ENT, PRC and posterior portions of EVC, with the left ENT/PRC cluster extending to abut the anterior medial hippocampus and the EVC cluster extending anteriorly along the lingual gyrus into the left posterior hippocampus. Where the activity involved the ENT and abutted the hippocampus, the location bordered the pre/parasubiculum much more anteriorly than that for scene construction. Therefore, naturalistic 3D scene representations may engage a circumscribed portion of the anterior pre/parasubiculum in unison with the PHC, RSC and PCC. By contrast, more general or abstracted forms of spatial imagery, such as objects on a 2D space (see also Constantinescu et al., 2016), might recruit a more anterior portion of ENT abutting the very anterior pre/parasubiculum. The different parts of the hippocampus and distinct cortical regions engaged by scenes and arrays, despite the close matching of the tasks, precludes the view that scenes are merely being enabled by processing sets of associations similar to those underpinning array construction. What we document here are separable mental construction processes giving rise to distinct types of representation in and around the hippocampus.

Considering the posterior hippocampal activation during array construction, this area has been implicated in a broad range of cognitive processes (Strange et al., 2014; Poppenk et al., 2013; Zeidman and Maguire, 2016), including spatial memory (Maguire et al., 2006; Moser and Moser, 1998) and mnemonic processing of items in a 2D space (de Rover et al., 2011). While our results reflect involvement of the posterior hippocampus in mental imagery of objects in a 2D rather than 3D space, it is unlikely that the posterior hippocampus is only involved in this form of mental imagery. The anatomy of the most posterior portion of the human hippocampus is particularly complex (Dalton et al., 2017), with much still to learn. Ultra-high resolution MRI investigations at the level of subfields are required to further inform our understanding of posterior hippocampal contributions to specific cognitive processes.

The Construct Objects task was not designed to be a close match for the array and scene tasks, but was included to inform about the brain areas engaged during object construction and object only associations, where spatial context was irrelevant. Of note, vividness of the imagery and memory for the objects in this task was significantly better than those in the array and scene construction tasks. Therefore, any fMRI results should be interpreted with this is mind. PRC recruitment during object imagery would be predicted, and indeed was found, considering the strong association between the PRC and object-centred cognition (Murray et al., 2007). Overall, the object task engaged very similar areas to those recruited for array construction, namely, the PRC, ENT abutting the very anterior medial left hippocampus. This may reflect a generic area for non-scene based associative processing. Where object construction differed from both array and scene tasks was in the activation of the right anterior lateral hippocampus in the region that aligns with the location of the prosubiculum/CA1. This finding is concordant with the prediction of Dalton and Maguire (2017), based on neuroanatomical considerations, where PRC, ENT and prosubiculum/CA1 are heavily interconnected (Li et al., 2017; Insausti and Munoz, 2001). Therefore, as with the array and scene construction tasks, the mental construction of isolated objects engaged a differentiable portion of the hippocampus.

Our results show that for associations between objects, between objects and 2D space or objects and 3D space, the hippocampus does not seem to favour one type of representation over another; it is not a story of exclusivity. Rather, there may be different circuits within the hippocampus, each associated with different cortical inputs, which become engaged depending on the nature of the stimuli and the task at hand. This may explain why it has been so difficult to reconcile hippocampal theories that have generally focused on one type of process or representation. Our results may also explain why disparate patterns of cognitive preservation and impairment are reported in patients with bilateral hippocampal lesions. For any individual, damage may affect particular subregions within the hippocampus more than others. These subtle case-by-case differences in the microscopic topography of damage and sparing may impact on cognition in different ways that as yet remain undetectable by current MRI technology.

While some theoretical accounts have posited that distinct areas within the MTL may preferentially process specific types of representation (Graham et al., 2010; Barense et al., 2005), perhaps surprisingly, such representational differences have not typically been extended to processing *within* the hippocampus. Non-human primate tract tracing studies have shown clear differences in how different cortical and subcortical brain regions interact not only with specific hippocampal subfields (Ding et al., 2000; Rockland and Van Hoesen, 1999; DeVito, 1980) but also disproportionately interact with specific portions of subfields along the longitudinal (anterior–posterior) and transverse (distal–proximal) axes of the hippocampus (Insausti and Munoz, 2001). In recent years, functional differentiation down the long axis of the hippocampus (reviewed in Poppenk et al., 2013; Nadel et al., 2013; Strange et al., 2014; Zeidman and Maguire, 2016) and subfield-specific hypotheses (Guzman et al., 2016; Berron et al., 2017; Libby et al., 2012) have received increasing attention. Our results further emphasise the importance of considering the hippocampus as a heterogeneous structure, and that a focus on characterising how specific portions of the hippocampus interact with other brain regions may promote a better understanding of its role in cognition.

It remains possible that other factors may have affected our results. For example, it could be that participants engaged in more size transformation of objects, or visualisation of objects in a more distant space, during the Construct Scene task. We are, however, unaware of any evidence for MTL involvement in these processes (Larsen et al., 2000). Future work is needed to precisely characterise the different information processing streams within the human hippocampus, both anatomically and functionally. Presumably these circuits are linked, but how and to what extent will also be important questions to address. In humans, little is known about intrahippocampal functional connectivity or even connectivity between specific hippocampal subfields and the rest of the brain. Use of ultra-high resolution (f)MRI is clearly warranted to help move beyond an ‘either/or’ view of the hippocampus to a more nuanced understanding of its multifaceted contributions to cognition.

## Acknowledgements

This work was supported by a Wellcome Principal Research Fellowship to E.A.M. (101759/Z/13/Z) and the Centre by a Centre Award from Wellcome (203147/Z/16/Z). The authors are grateful to David Bradbury for technical assistance.

## Abbreviations

ENT: Entorhinal cortex
EVC: Early visual cortices
PCC: Posterior cingulate cortex
PHC: Posterior parahippocampal cortex
PRC: Perirhinal cortex
RSC: Retrosplenial cortex

